# A PERK-FOXO1 AXIS LINKS DNA DAMAGE TO FIBROBLAST SURVIVAL IN DIFFUSE CUTANEOUS SYSTEMIC SCLEROSIS

**DOI:** 10.64898/2026.02.17.706443

**Authors:** Lamia Khan, Junquin Wang, Carmanah Hunter, Charmaine van Eeden, Desiree Redmond, Lisa Willis, Caylib Durand, Jan Storek, Kareem Jamani, Udo Mulder, Murray Baron, Janet Pope, Elena Netchiporouk, Jan Willem Cohen Tervaert, Harissios Vliagoftis, Robert Gniadecki, Mohammed Osman

## Abstract

**Objective:** Diffuse cutaneous systemic sclerosis (dcSSc) is a life-limiting fibrotic disease. We and others have shown that dcSSc fibroblasts accumulate numerous somatic mutations associated with senescence-like features; however, the mechanism(s) enabling their survival remain unclear.

**Methods:** Skin biopsies were obtained from lesional tissues from dcSSc (n=10), dcSSc treated with autologous hematopoietic stem cell transplantation (ASCT, n=8) or 7 age/sex-matched healthy controls. Primary dermal fibroblasts were generated from biopsies. Spatial RNA sequencing, immunoblotting, confocal microscopy, and functional assays were used to mechanistically delineate signaling pathways linking DNA-damage with fibroblast survival.

**Results:** dcSSc fibroblasts demonstrated increased pH2AX DNA double-strand-break foci yet remained apoptosis resistant. These cells displayed features of metabolic-stress remodeling, including mitochondrial hyperpolarization, increased reactive oxygen species production, and enhanced mitochondrial biogenesis. Spatial transcriptomics and subsequent biochemical analyses identified activation of a PERK/ATF4/FOXO1 axis, characterized by PERK phosphorylation, selective ATF4 translation, FOXO1 nuclear translocation, and induction of downstream antioxidant and metabolic programs. In contrast, fibroblasts from post-ASCT patients exhibited normalization of DNA-damage markers and mitochondrial parameters without ATF4/FOXO1 activation. Pharmacologic inhibition of either PERK or FOXO1 selectively restored mitochondrial-dependent apoptosis in dcSSc fibroblasts, demonstrating that this axis is required for their survival following extensive genomic injury.

**Conclusion:** dcSSc fibroblasts persist despite substantial genomic injury by engaging a PERK/ATF4/FOXO1 metabolic-adaptation program that suppresses mitochondrial-dependent apoptosis. This survival axis is not present after ASCT. Targeting PERK or FOXO1 restores apoptosis selectively in dcSSc fibroblasts, highlighting its potential use as a therapeutic target for eliminating pathogenic senescence-like fibroblasts in dcSSc.

**Highlights:**

- Both *ex-vivo* skin and *in-vitro* primary dermal fibroblasts derived from dcSSc patients have a higher frequency of intrinsic DNA damage signals and senescence-associated features; yet they evade mitochondrial-dependent apoptosis.
- Pathogenic dcSSc fibroblasts rewire their metabolism, characterized by mitochondrial hyperpolarization and elevated ROS.
- Spatial transcriptomics and functional analyses reveal a PERK/ATF4/FOXO1 stress-adaptation axis that drives fibroblast survival in dcSSc.
- This maladaptive survival program characterized by increased genotoxic stress, and mitochondrial remodelling is absent in post-ASCT fibroblasts.
- Targeting PERK or FOXO1 selectively sensitizes dcSSc fibroblasts to apoptosis revealing a potential promising therapeutic strategy in dcSSc.

**Graphical abstract:** 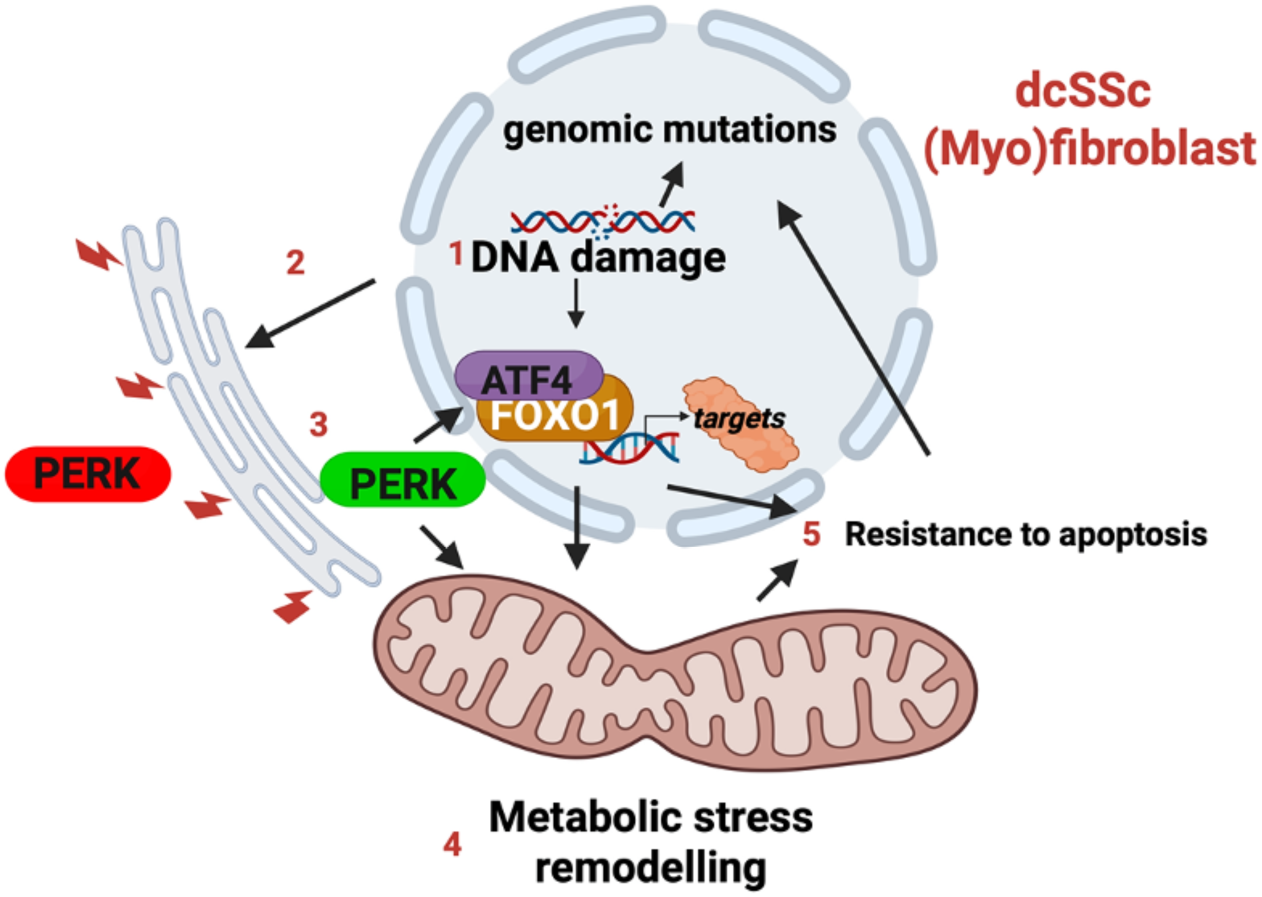

## Introduction

Diffuse cutaneous systemic sclerosis (dcSSc) is an autoimmune connective tissue disorder dominated by progressive, widespread fibrosis of the skin and internal organs, associated with vascular dysfunction and persistent immune activation ^1^. Despite therapeutic advances, the five-year mortality rate remains high, and patients continue to experience poor outcomes ^2,3^. The most effective treatment for dcSSc is autologous stem cell transplantation (ASCT), which offers greater improvement in skin fibrosis than other therapies ^4,5^. In dcSSc, fibrosis is promoted by activated fibroblasts (FB) which adapt a myofibroblast phenotype ^6^, which is detectable both *in vitro* and *in vivo* – suggesting that various epigenetic adaptations may be present in dcSSc FB independently of inflammatory mediators derived from immune cells ^6^. This observation may explain the lack of spontaneous regression of fibrosis in many dcSSc patients despite immunosuppression ^3^.

Several groups have highlighted metabolic remodelling and resistance to apoptosis as important molecular adaptations in dcSSc FB ^7,8^. These changes are linked to markers of cellular senescence (e.g., increased expression of p16 and p21) and the secretion of profibrotic cytokines (e.g., IL-6, IL-8, or TGF-β1), which may collectively amplify signals that lead to increased skin inflammation, fibrosis, evasion of intrinsic apoptosis, and a feed-forward sustained senescence-like signature ^6,8–10^. Recognizing that somatic mutation accumulation is a known driver of cellular senescence phenotype, we performed whole-exome sequencing of fibrotic dcSSc skin to investigate whether genomic alterations are implicated in this dysregulated myofibroblast phenotype ^11^. We found that dermal FB are hypermutated, harboring numerous somatic mutations and often included recurrent changes in cancer driver genes involved in DNA repair, chromatin regulation, and genome integrity. These mutations exhibited a mutational signature typical of age- and replication-associated “clock-like” processes. Those findings, which were subsequently confirmed independently ^12^, suggest that dcSSc FB may engage survival and stress-adaptation programs. Consistent with this idea, our spatial RNA sequencing analysis of SSc dermis revealed that FB-enriched regions displayed prominent mitochondrial perturbations pointing to a link between genotoxic signaling and metabolic dysregulation ^13^. As a result, we hypothesized that dcSSc FB acquire maladaptive stress responses similar to those implicated in fibrotic remodeling of the heart and liver ^14–16^ that allow them to withstand genomic injury by evading mitochondrial-dependent apoptosis; and ASCT may normalize these mechanisms in FB.

An important pathway involved in this maladaptive stress response is activation of protein kinase RNA-like endoplasmic-reticulum kinase (PERK) and downstream activation of its targets activating transcription factor 4 (ATF4) ^17^ and forkhead box 1 (FOXO1) ^18,19^. The PERK/ATF4/FOXO1 axis is an essential component of the integrated stress response that connects endoplasmic-reticulum homeostasis and mitochondrial metabolism to cell-fate decisions ^17,20^. Activation of PERK in response to genotoxic stress transiently suppresses protein synthesis while promoting selective translation of ATF4, which coordinates adaptive responses through autophagy, antioxidant defense, and amino-acid metabolism ^17,20^. PERK also activates the transcription factor FOXO1 ^18^, which translocates to the nucleus to induce transcription of genes that promote cell survival ^21^.

In this mechanistic study, we postulate that dcSSc fibroblasts evade apoptosis via a PERK/ATF4/FOXO1-dependent pathway which enables them to withstand extensive genotoxic stress. Importantly, inhibition of PERK or FOXO1 restored apoptosis, revealing a targetable mechanism with therapeutic implications in dcSSc (*see graphical abstract*).

## Materials and Methods

### Patients, clinical information, skin samples and storage

All patients met the 2013 European Alliance of Associations for Rheumatology/American College of Rheumatology (ACR/EULAR) classification criteria for dcSSc ^22^. This work was approved by the University of Alberta Research Ethics Board (Pro00085583, Pro00090050) which includes members of the public. We recruited 10 dcSSc patients and 8 dcSSc who were treated with ASCT (post-ASCT, median follow-up period 12 months) who were not receiving any immunomodulatory therapy at the time of sample collection from the University of Alberta and Calgary. We excluded patients with a known history of a malignancy, age >65y, or with a disease duration >5y from the first non-Raynaud’s symptom. We obtained all our patient’s clinical history/information including presence or absence of anti-RNA polymerase III antibodies, anti-centromere antibodies (ACA), anti-topoisomerase I antibodies (anti-ATA/Scl70), and interstitial lung disease (ILD). Four-millimeter skin punch biopsies were obtained from the distal forearm (∼5 cm from the ulnar styloid), as we previously described ^11^. A 1 mm section of each biopsy was used to generate primary dermal FB, while the remainder was embedded in optimum cutting temperature (OCT) medium and stored at –80°C. Control material was obtained from 7 healthy individuals (healthy control or HC) of a similar median age as the dcSSc patients (detailed clinical characteristics in **supplementary table 1**).

### Antibodies and reagents

Antibodies (with vendors and catalogue numbers) utilized in our experiments included: pH2AX (Cell Signaling Technology, cat# 80312S), Total H2AX (Cell Signaling Technology, cat# 7631S), p21 (Cell Signaling Technology, cat# 2947S), p16 (Cell Signaling Technology, cat# 80772S), alpha-smooth muscle actin (Abcam, cat#ab7817), ß-actin (Cell Signaling Technology, cat# 8457S), cleaved caspase 3 (Cell Signaling Technology, cat# 9661S), cleaved caspase 9 (Cell Signaling Technology, cat# 20750S), OPA1 (Cell Signaling Technology, cat# 67589S), DRP1 (Cell Signaling Technology, cat# 8570S), p-DRP1 (Abcam, cat# ab193216), tubulin (Abcam, cat# ab6160), PERK (Cell Signaling Technology, cat# 3192S), p-PERK (ThermoFisher scientific, cat# PA5-102853), FOXO1 (Cell Signaling Technology, cat# 2880S), p-FOXO1 (Cell Signaling Technology, cat# 9461S), MFN1 (Cell Signaling Technology, cat# 14739S), MFN2 (Cell Signaling Technology, cat# 9482S), ATF4 (Cell Signaling Technology, cat# 11815T).

### Cell culture and treatments with pharmacological agents

We generated primary dermal FB from punch biopsies obtained from study patients and controls, as we previously described ^11^. FB were phenotyped using immunofluorescence microscopy confirming the presence of FB-specific markers ^11^. All primary FB were maintained in DMEM containing 1% penicillin-streptomycin and 10% FBS (ThermoFisher scientific,) in 5% CO2 at 37°C in a humidified environment. We regularly performed mycoplasma test (MycoAlert™ Mycoplasma Detection Kits, Lonza) on FB to detect any potential contamination. All *in vitro* experiments were performed from low passage dermal FB (passage 2-5), at a confluency of 70-80%, then grown in DMEM containing 0.1-0.2% FBS for 18-20 hours (for synchronization) prior to specific experiments. To inhibit PERK kinase activity, we used PERK inhibitor (GSK 2606414, 1µM) (TOCRIS Biotechne, cat#5107) for 2 hours. Similarly, we treated FB with a FOXO1 inhibitor (AS1842856, 1µM) (Sigma Aldrich, cat# 344355) for 4-6 hours. To induce cell death, we treated cells with 4-<u>H</u>ydroperoxy <u>c</u>yclophosphamide (4-HC, 2 µM) (Santa Cruz Biotechnology, cat#sc-206885) for 23 hours, then measured apoptosis. To induce DNA damage, we treated cells with etoposide (Sigma Aldrich, cat# E1383, 50 µM) for 2 hours, then measured indicators associated with DSBs (phosphorylated histone 2AX or pH2AX), phosphoSer256-FOXO1 (normalized to total FOXO1) or ATF4 levels via immunoblot.

### Immunoblot (IB) and co-immunoprecipitation

FB cells were grown as above then subjected to SDS-PAGE/IB analysis. Briefly, cells were lysed using radio-immunoprecipitation Assay (RIPA) buffer (ThermoFisher scientific, cat# 89900) mix containing 1mM phenylmethylsulfonyl fluoride (PMSF), 1X protease inhibitor (ThermoFisher scientific, cat#78429), 1X phosphatase inhibitor (ThermoFisher scientific, cat#78420). Protein quantification in each cell lysate was measured via a Micro BCA™ Protein Assay Kit (ThermoFisher scientific, cat# 23235). Proteins were resolved using SDS-PAGE (in a 8 or 12% gel acrylamide: bis-acrylamide (fisher scientific, cat#HBGR337500)), transferred to a 0.2 micron nitrocellulose membrane, blocked in 5% BSA in tris-buffered saline (TBS) containing 0.1% Tween 20) then incubated overnight at 4°C using the indicated primary antibodies which were subsequently detected, using fluorescently-labeled secondary antibodies (goat anti-rabbit IR Dye 800 (Li-Cor, cat# 926-32211), goat anti-mouse IR Dye 800 (Li-Cor, cat# 926-32210), or Donkey anti-rabbit IR Dye (Li-Cor, cat#680 926-68073), as appropriate) then visualized using a Li-Cor Odyssey CLx imaging system. IB from each FB group was repeated with at least 2 independent replicates. For co-immunoprecipitations, cells were harvested then washes with ice-cold PBS and lysed in PBS containing 0.1 % Triton X-100 in DPBS with 1 mM PMSF and benzonase for 30 min on ice. 100 µg of cell lysate was incubated with the immunoprecipitation antibody (or IgG control) overnight at 4°C, with rotation, then protein G beads were added and incubated for 4 hrs at 4°C with rotation. Proteins were eluted with SDS-PAGE loading dye then subjected to SDS-PAGE/IB.

### Quantitative RT-PCR (qRT-PCR)

Total RNA from each FB group was extracted using a Qiagen RNeasy Mini Kit (cat# 74104). Subsequently, qRT-PCR was performed via a TaqMan™ Fast Virus 1-Step Master Mix (ThermoFisher Scientific, cat# 4444436) and specific TaqMan™ primers; then samples were analyzed using a QuantStudio 3 Real Time PCR system (Applied Biosystems). mRNA transcript levels were normalized to GAPDH. The relative gene expression was evaluated by plotting it via a 2^−ΔΔCT^. The Taqman probes (Thermo Fisher Scientific) used for qRT-PCR (with targets) included: Hs02596861_s1 (D-loop), Hs02596876_g1 (ND4), Hs02596867_s1 (CytB), Hs00231106_m1 (FOXO1), Hs02758991_g1 (GAPDH), Hs00188166_m1 (Succinate dehydrogenase or SDHA), Hs00167309_m1 (SOD2), Hs01595220_g1 (COX7C), Hs00176875_m1 (PDK4), Hs01561847_m1 (PDK1), Hs00176865_m1 (PDK2).

### Immunofluorescence (IF) staining

For dermal staining, 10 μm skin sections stored in -80°C were mounted on glass slides and then stored at -80°C as described before ^23^. Prior to staining, sections were brought to room temperature, then permeabilized with 0.1% Triton X-100 (Sigma-Aldrich, cat# T9284), and blocked with 2% bovine serum albumin (BSA). phospho-H2AX (pH2AX) levels were determined using mouse anti-pH2AX (Cell Signaling Technology, cat# 80312), then stained with rabbit anti-vimentin to identify FB (Cell Signaling Technology, cat# 5741). Sections were counterstained with donkey anti-mouse Alexa Fluor 647 (ThermoFisher Scientific, cat# A-31571) and anti-rabbit Alexa Fluor 555 (ThermoFisher Scientific, cat# A-31572) secondary antibodies. Nuclei were stained with 4’,6-diamidino-2-phenylindole (DAPI, ThermoFisher Scientific, cat# 62248), and visualized using an Olympus IX-8I spinning disk confocal microscope. For FB IF imaging, cells were directly grown (70-80 % confluence) and treated (as indicated), then fixed with 4% PFA, permeabilized with 0.1% Triton X-100, then stained with the specific primary antibodies overnight at 4°C, and then counterstained with appropriate secondary antibodies and DAPI, then mounted in ProLong Gold antifade and imaged as before. ImageJ software was used to quantify the signal intensity.

### Mitochondrial network analysis

Dermal FB directly grown on glass-bottom confocal dishes (VWR, cat#75856-740) to 70-80 % confluence, then grown at 0.1-0.2% FBS in DMEM as before. To label mitochondria, the cells were incubated with 200 nM MitoTracker™ Red CMXRos (ThermoFisher Scientific, cat# M7512) for 30 minutes in a 37°C CO₂ incubator, fixed with 4% PFA then permeabilized using 0.1% Triton X-100 and nuclei were counterstained with DAPI. To visualize mitochondria, z-stack images were captured using an Olympus IX-81 spinning disk confocal microscope with a 60X oil immersion objective. For mitochondrial network analysis, FIJI with the Mitochondria Analyzer plugin (open-source image processing software) was used to analyze 20 cells per high powered field in at least 5 distinct visual fields per confocal dish for each subject.

### Mitochondrial membrane potential determination (Ψ_mt_) and reactive oxygen species (ROS) detection

To detect changes in mitochondrial membrane potential (Δ Ψ_mt_), present in FB, cells from each group (HC, dcSSc, post-ASCT) were grown as before then incubated using the cationic dye tetramethylrhodamine (TMRM, (ThermoFisher, cat#M20036)) for 30 minutes at 37°C in a CO₂ incubator, then analyzed using flow cytometry (Attune NxT Flow Cytometer (ThermoFisher)). To establish the background levels of TMRM fluorescence, we treated cells from each group with carbonyl cyanide 3-chlorophenylhydrazone (CCCP), then gated on TMRM^pos^ cells. TMRM mean fluorescence intensity (MFI) was quantified using the FlowJo software. To detect ROS levels, FB from each group were grown as before to 70-80 % confluence, then ROS was quantified using flow cytometry (Attune NxT) by measuring CellROX™ Green fluorescence intensity (ThermoFisher Scientific, cat#C10444) with background levels normalized to unstained cells, and the fluorescence intensity was quantified using FlowJo software (BD Biosciences) version 10.9.0.

### Apoptotic cell death detection

Apoptotic cell death was detected using terminal U-nicked end labelling (TUNEL, Sigma Aldrich, cat# s7110) following the manufacturer’s instructions. Briefly, cells were seeded on coverslip and treated with 2µM 4-HC for 23 hours. Then cells were fixed and stained with the TUNEL assay reagents, and intra-nuclear. TUNEL positive cells were visualized with widefield fluorescence microscopy. Alternatively, cells were treated with 2 µM 4-HC, a PERK inhibitor (1 µM), or a FOXO1 inhibitor (1 µM), then cleaved caspase 3 and cleaved caspase 9 were quantified using IB.

### Spatial mRNA sequencing

We re-analyzed spatial RNA sequences that we recently reported using the Visium spatial RNA (10X Genomics, Pleasanton, CA, USA), and 10 μm SSc patient-derived dermal sections^13^. We aligned sequences obtained to the hg38 reference genome using Space Ranger v2.1.0 and FB clusters were augmented by obtaining dermal FB obtained from single cell RNA sequencing which was obtained from an independent study ^24^. Dimensionality reduction and visualization were performed using UMAP via Seurat v5.1. We created heatmaps for differential expression in the fibroblast clusters (Clusters 0 and 1) using GO and KEGG pathway analysis via ClusterProfiler v3.19; and in R Studio v4.4.1 to identify targets predicted to be downstream of the unfolded protein response or FOXO1.

### Statistical analysis

All FB results are shown as the mean ± SEM of independent biological replicates. All statistical analyses were conducted using GraphPad Prism 10 software. Data normality was tested using the Shapiro-Wilk normality test. If data were normally distributed, a parametric test was used. If not, a non-parametric analysis was performed. To assess differences between two groups, either an unpaired or paired Student’s t-test was applied, as appropriate. For non-parametric comparisons, the Mann–Whitney U test was used for unpaired data, while the Wilcoxon matched-pairs signed-rank test was applied for paired non-parametric analyses. For paired parametric data, a ratio-paired t-test was used. For comparisons involving more than two groups, a one-way or two-way analysis of variance (ANOVA) was used for parametric data, while the Kruskal-Wallis test was applied for non-parametric analyses, followed by a post-hoc test.

## Results

### Double-stranded DNA breaks (DSB) are more abundant in dcSSc FB

We and others recently showed that FB from patients with dcSSc have increased genomic mutations ^11,12^. Since mutations result from genomic instability and DNA damage (most commonly DSB), we quantified DSB in dermal sections and cultured FB from dcSSc patients before and after therapeutic ASCT, comparing them with the age and sex-matched healthy controls. DSBs were assessed by enumerating pH2AX foci ^25,26^.

Dermal sections from dcSSc demonstrated a higher frequency of nuclear pH2AX DSB damage foci in vimentin-positive FB than either post-ASCT or HC samples (**Fig. 1A, Supplementary fig. 1A**). Also, dermal FB from dcSSc showed more pH2AX DSB damage foci than FB from HC or post-ASCT patients (**Fig. 1B**). Consistent with those findings, the total pH2AX levels (normalized to total intracellular H2AX) was significantly higher in dcSSc FB, compared to both HC and post-ASCT groups (**Fig. 1C**). Given that DNA damage is known to promote pro-fibrotic myofibroblast activation, we next assessed fibroblast activation across groups using a marker, alpha-smooth muscle actin (α-SMA). The α-SMA expression was significantly higher in dcSSc FB compared to HC and normalized in post-ASCT (**Supplementary fig. 1B**), suggesting more active myofibroblast in dcSSc. Collectively, our data suggests that activated dcSSc FB harbor the highest DSB burden among all groups, whereas ASCT is associated with reduced FB activation and genomic DNA damage.

**Fig. 1.**
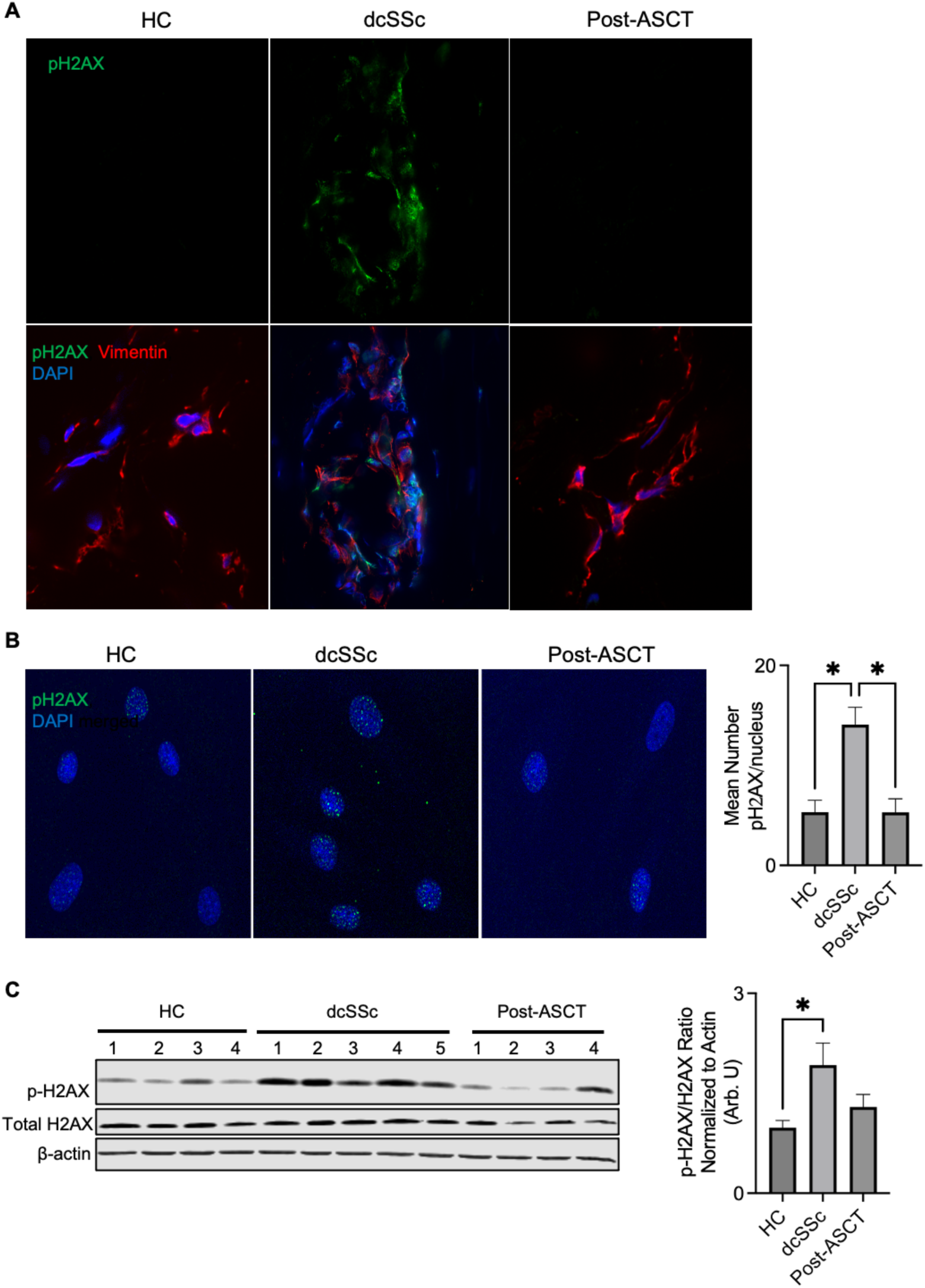
Double stranded DNA breaks are more abundant in dcSSc FB. **A)** Representative confocal immunofluorescence images of pH2AX (green) and DAPI-stained nuclei (blue) at 60X magnification in skin section from independent biological replicates of HC (n=5), dcSSc (n=7), and post-ASCT (n=4) to analyze the damaged DNA foci. **B)** Representative confocal immunofluorescence images of pH2AX (green) and DAPI-stained nuclei (blue) at 40X magnification in independent biological replicates of primary dermal fibroblasts (FB) from HC (n=6), dcSSc (n=7), and post-ASCT (n=4) with enumeration nuclear DNA damage foci. Confocal images were quantified using ImageJ software. **C)** Immunoblot analysis of pH2AX /total H2AX ratio (normalized to beta-actin) in independent biological replicates of HC (n=6), dcSSc (n=7), and post-ASCT (n=6). All represent the means ± SEM. Statistical significance was determined by one -way ANOVA. Statistical significance was determined when *p<0.05, **p<0.005 and ***p<0.0005 using GraphPad Prism.

### dcSSc fibroblasts have senescence-like signatures and are more resistant to apoptosis

Persistence of DSB is known to induce cellular senescence characterized by p21 and p16 – mediated cell cycle arrest ^27,28^ ^29^ ^30^, reduced growth ^31^ and resistance to mitochondrial-dependent apoptosis ^8,32,33^. Firstly, we confirmed that dcSSc FB showed a strong senescence-like phenotype characterized by increased p16 (**Fig. 2A**) and p21 levels (**Supplementary fig. 2A**). Consistent with previous reports ^9^, p16 levels were more elevated in dcSSc compared with HC and post-ASCT FB. Moreover, growth rate was significantly reduced in dcSSc FB compared to both HC and post-ASCT groups (**Supplementary fig. 2B**), further supporting senescence-like signature in dcSSc FB.

**Fig. 2.**
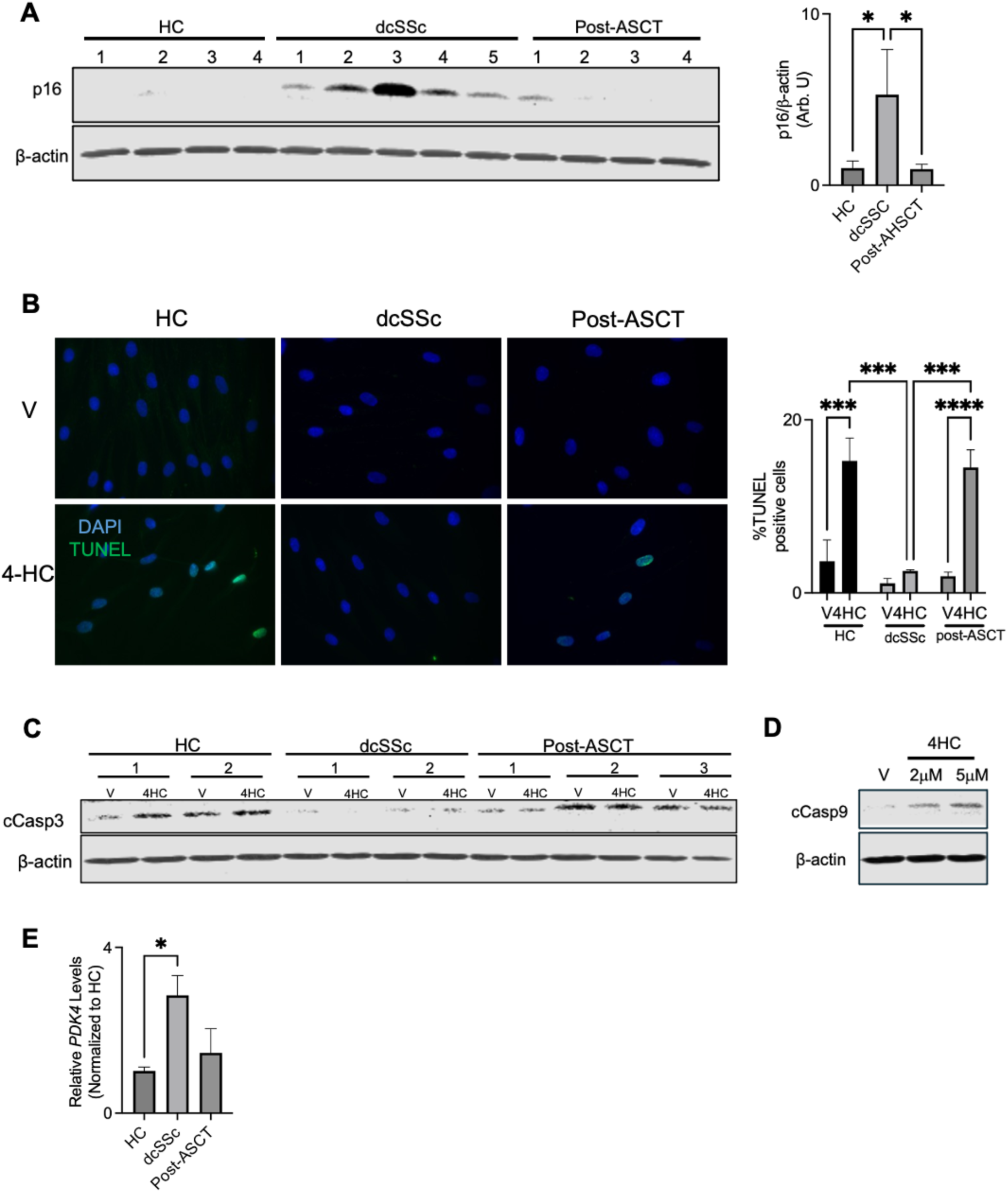
dcSSc FB have increased senescence-like signatures and resistance to apoptosis. Primary dermal FB from independent biological replicates of HC, dcSSc, or post-ASCT patients were used to determine the levels of **A)** p16 (N: HC=7, dcSSc=8 and post-ASCT=8) with relative abundance normalized to β-actin. **B)** TUNEL/Immunofluorescernce were used to detect the frequency of apoptosis (N, HC=3, dcSSc=4 and post-ASCT=4 X 2 independent biological replicates); or **C)** Cleaved Caspase 3 (normalized to β-actin as a loading control) from HC, dcSSc, or post-ASCT FB (N, HC=3, dcSSc=3 and post-ASCT=4 X 2 independent biological replicates) with representative immunoblot at baseline or in the presence of 4-HC. **D)** HC FB were treated with 4-HC then cleaved Caspase 9 levels were detected to show that apoptosis is mitochondrial dependent. **E**) *Pyruvate dehydrogenase kinase 4* (*PDK4*) (N: HC=5, dcSSc=7and post-ASCT=5, independent biological replicates) are elevated in dcSSc but not HC or post-ASCT FB - suggesting that resistance to apoptosis may be driven via stress-associated mitochondrial remodeling. *PDK4* levels were directly measured in HC, dcSSc, or post-ASCT primary FB using qRT-PCR with results normalized to relative *PDK4* mRNA in HC. All data represent the means ± SEM. Statistical significance was determined by one-way and two-way ANOVA as appropriate. GraphPad Prism determined statistical significance when *p<0.05, **p<0.005 and ***p<0.0005.

Secondly, we induced mitochondrial-dependent apoptosis using 4-HC (the active ingredient of cyclophosphamide) ^34^ then quantified apoptosis via TUNEL/IF and confirmed that dcSSc FB were highly resistant to apoptosis (**Fig. 2B**). Thirdly also measured cleaved caspase 9 (cCasp9), which is associated with mitochondrial-dependent intrinsic apoptosis pathway, and cleaved caspase 3 (cCasp 3, downstream of cCasp9). We showed that cCasp3 was reduced in dcSSc FB compared to post-ASCT and HC FB (**Fig. 2C**) - suggesting that dcSSc harboured more mitochondrial-dependent apoptosis resistance, particularly as 4-HC induced a mitochondrial-dependent apoptosis in HC as suggested by increased levels of both cleaved cCasp 9 and 3 (**Fig. 2C, D**). Finally, dcSSc had increased levels of pyruvate dehydrogenase kinase 4 (*PDK4),* which promotes glycolytic metabolic stress reprogramming ^35^ (**Fig. 2E**), but not other PDKs (*PDK1* or *PDK2)* (**Supplementary fig. 2C, D**) *-* suggesting that resistance to apoptosis may be associated with metabolic stress remodeling in dcSSc.

### dcSSc develop molecular changes consistent with metabolic stress remodeling

Resistance to mitochondrial-dependent apoptosis is promoted by metabolic adaptations that limit the release of pro-apoptotic mediators. These metabolic adaptations are characterized by elevated mitochondrial membrane potential (ΔΨmt) ^36^. It is also associated with increased production of reactive oxygen species (ROS) at ETC complexes I (ND4) and III (cytochrome b), and mitochondrial biogenesis (referred to as metabolic-stress remodeling) ^37–39,40,41^. To determine this, we first measured ΔΨ_mt_ and found it markedly increased in dcSSc FB compared to HC, with normalization to HC levels in post-ASCT FB (**Fig. 3A**). Then, we measured the mRNA expression of mitochondrial ETC complexes ^13^. dcSSc FB had increased mRNA levels of both *ND4* (ETC Complex I) and *Cytochrome b* (ETC Complex III), whereas transcripts from ETC complexes less associated with ROS generation (e.g. *succinate dehydrogenase A SDHA* (complex II) or *Cox7C* (Complex IV) were not increased relative to HC or post-ASCT FB (**Fig. 3B**). These transcriptional changes were functionally relevant, as dcSSc FB displayed significantly higher levels of intracellular ROS than both comparator groups (**Fig. 3C**). Finally, given that chronic mitochondrial stress remodeling may promote mitochondrial biogenesis, we quantified the mitochondrial D-loop as a marker of newly synthesized mtDNA and observed increased levels in dcSSc FB (**Fig. 3D**). Together, these findings indicate that dcSSc FB undergo metabolic-stress remodeling characterized by mitochondrial hyperpolarization, enhanced ROS production, and increased mitochondrial biogenesis.

**Fig. 3.**
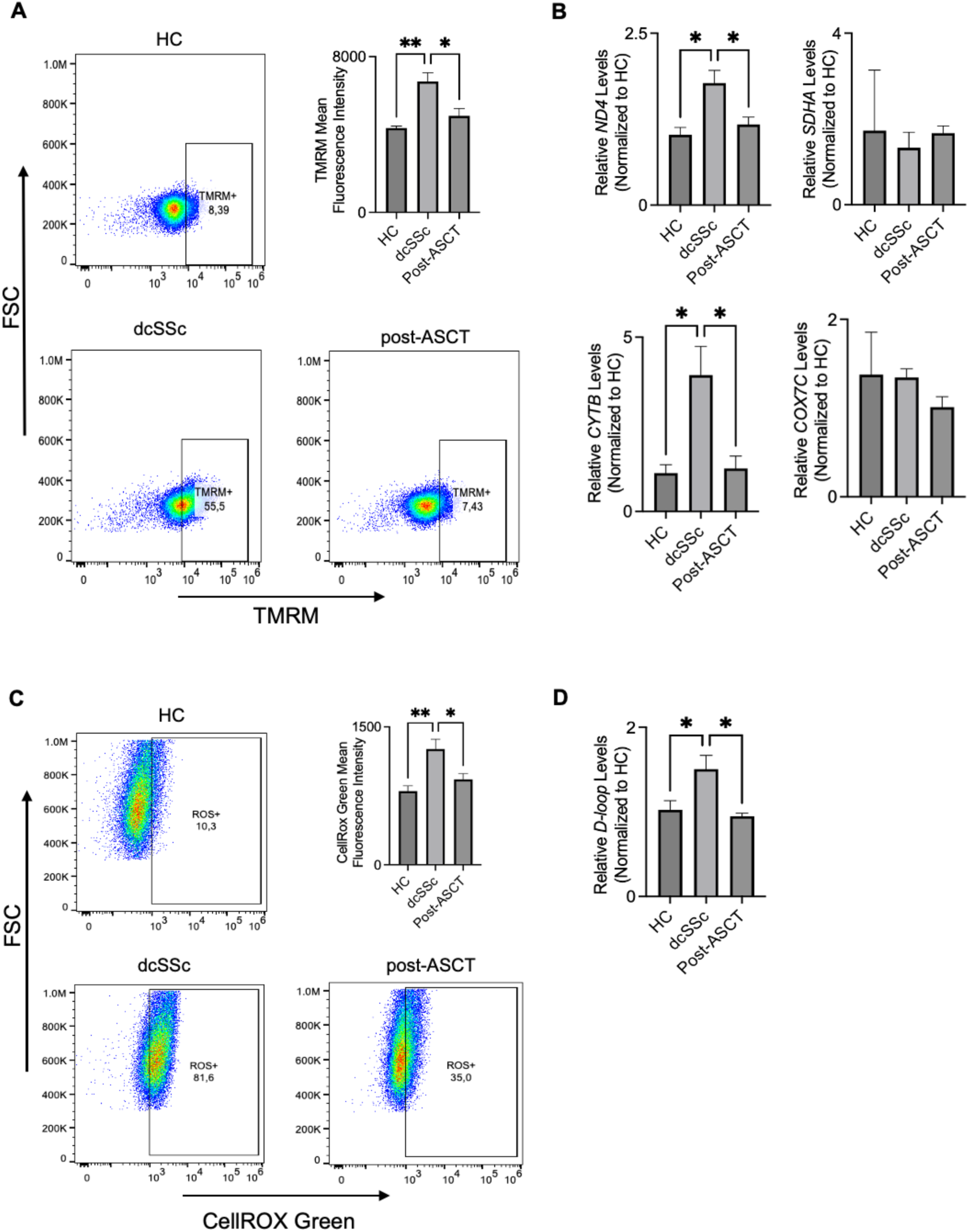
dcSSc FB have increased ΔΨmt and ROS with associated metabolic stress. remodeling. **A)** Primary dermal fibroblasts (FB) were used to determine the resting mitochondrial membrane potential (ΔΨmt) using TMRM of live FB from independent biological replicates of each group, then quantified via flow cytometry analysis in HC (n=4), dcSSc (n=6), and post-ASCT (n=5) (with 2 independent replicates/group). **B)** qRT-PCR was used to determine the relative expression of various ETC complexes (*ND4,* complex I; *SDHA*, complex II; Cytochrome b (CytB, complex III), and Cyclo-oxygenase 7C (Cox7C, complex IV), N: HC=5, dcSSc=8 and post-ASCT=5, independent biological replicates). **C)** Total cellular ROS was determined by measuring the fluorescence intensity of CellROXgreen then quantified via flow cytometry from independent biological replicates of HC (n=6), dcSSc (n=6), and post-ASCT (n=5) FB (with two independent replicates/FB). **D)** Nascent mitochondrial DNA was quantified by quantifying D-loop levels using qRT-PCR (N: HC=5, dcSSc=6 and post-ASCT=5, independent biological replicates). All data were presented as the means ± SEM. Statistical significance was determined by one-way ANOVA. Using GraphPad Prism, statistical significance was determined when *p<0.05, **p<0.005 and ***p<0.0005.

### dcSSc FB have increased mitochondrial hyperfusion

One key adaptation supporting cell survival is mitochondrial hyperfusion^7^. Hyperfusion can be assessed morphologically (by quantifying the length and frequency of mitochondrial branches) and also biochemically by measuring the accumulation of the inner membrane fusion proteins, such as serine-637-phosphorylated dynamin-related protein 1 (pDRP1) ^18^, and the GTPase optic atrophy 1 (OPA1) ^42^, without reciprocal increases in outer-membrane fusion proteins (e.g. mitofusin 1 and 2 (MFN1, 2) ^9^.

We therefore evaluated mitochondrial architecture using MitoTracker Green, which fluorescently labels mitochondrial inner membranes. Confocal microscopy of MitoTracker Green-labelled cells revealed a striking increase in the number and length of mitochondrial branches in dcSSc FB compared with HC and post-ASCT cells (**Fig. 4A**). In addition, we detected elevated OPA1 and pDRP1 levels in dcSSc FB (**Fig. 4B, C)**, with no parallel increase in MFN1 and MFN2 levels (**Fig. 4D, E),** further documenting mitochondrial hyperfusion in dcSSc FB.

**Fig. 4.**
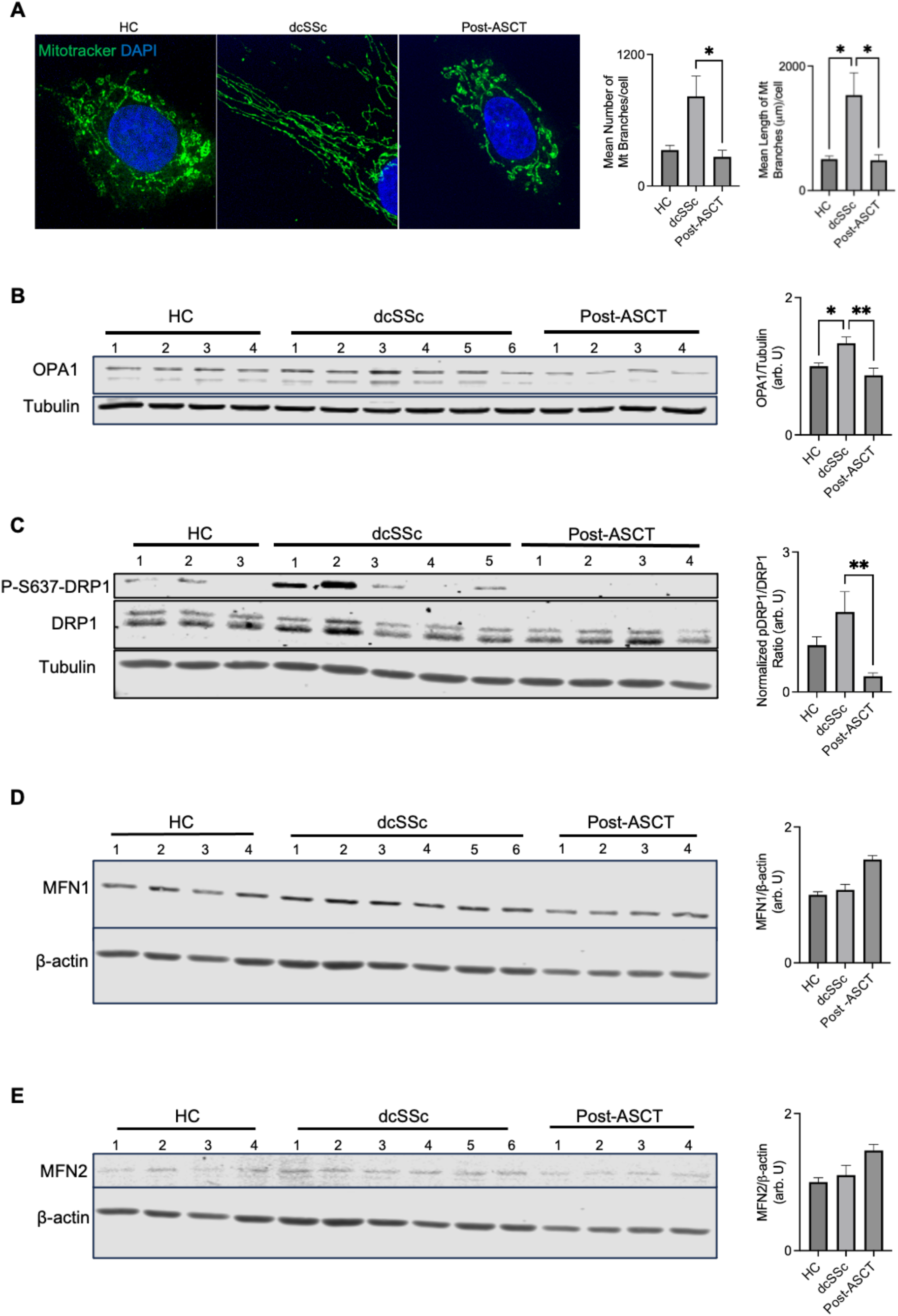
dcSSc FB develop stress-induced mitochondrial hyperfusion. We used primary dermal fibroblasts (FB) from HC, dcSSc, or post-ASCT to directly determine changes in mitochondrial morphology or biochemical changes associated with hyperfusion. **A)** Representative confocal immunofluorescence images using Mitotracker (green) and DAPI-stained nuclei (blue) at 60X magnification. We completed 2-3 independent technical replicates using biological replicates of HC (n=4), dcSSc (n=5), and post-ASCT (n=5) FB, then analyzed mitochondrial length, and branches using Fiji. **B)** We quantified the frequency of mitochondrial fusion proteins OPA1, **C)** ratio of p-DRP1/total DRP1, **D**) MFN1 or **E**) MFN2 in biological replicates of HC (n=7), dcSSc (n=8), and post-ASCT (n=8). All data were presented as the means ± SEM. Statistical significance was determined by one -way ANOVA. Statistical significance was determined when p<0.05 using GraphPad Prism.

### dcSSc FB have increased PERK, ATF4 and FOXO1 activation

PERK is a critical sensor that integrates genotoxic stress with metabolic stress remodeling ^15,20^. In settings where PERK promotes cell survival, it mediates mitochondrial hyperfusion ^43–45^. Activated PERK selectively enhances translation of ATF4, which in turn activates forkhead box 1 (FOXO1). Subsequently, FOXO1 becomes dephosphorylated at Ser256 and translocates to the nucleus, where it drives metabolic remodeling (e.g., increased PDK4) ^46^ and induces antioxidant enzymes such as *SOD2* ^21^. To assess whether the PERK/ATF4/FOXO1 axis is dysregulated in SSc, we re-analyzed our spatial RNAseq dataset in FB clusters ^47^, and noted that transcripts associated with stress response were increased in SSc skin (**Fig. 5A**) such as ATF4, FOXO1, antioxidants *SOD2* and *CAT*, and to a lesser extent *PDK4* (**Fig. 5A**, **Supplementary fig. 4**). We next directly measured PERK activation in cultured FB from the various groups. dcSSc FB had increased levels of phospho-PERK (pPERK), compared to HC, whereas post-ASCT FB showed modestly elevated pPERK, almost similar to dcSSc (**Fig. 5B**). Importantly, ATF4 levels were only markedly higher in dcSSc FB compared to both HC and post-ASCT FB (**Fig. 5C**) – suggesting that canonical PERK activation was only present in dcSSc, but not post-ASCT FB. Following its activation, ATF4 promotes *FOXO1* transcriptional activity ^48,49^. As a result, we directly quantified *FOXO1* mRNA levels in FB from our various groups by qRT-PCR. As predicted, dcSSc FB had markedly higher levels of *FOXO1* mRNA. PERK has also been shown to directly activate FOXO1 ^18^, which is determined via decreased Ser256 phosphorylation and increased nuclear translocation. To this end, dcSSc FB had decreased FOXO1 Ser256-phosphorylation, and increased nuclear FOXO1 levels (**Fig. 5D**-**5F**). This was functionally relevant as suggested by increased levels of the FOXO1 target *SOD2* (**Fig. 5G**) in dcSSc but not in HC or post-ASCT FB.

**Fig. 5.**
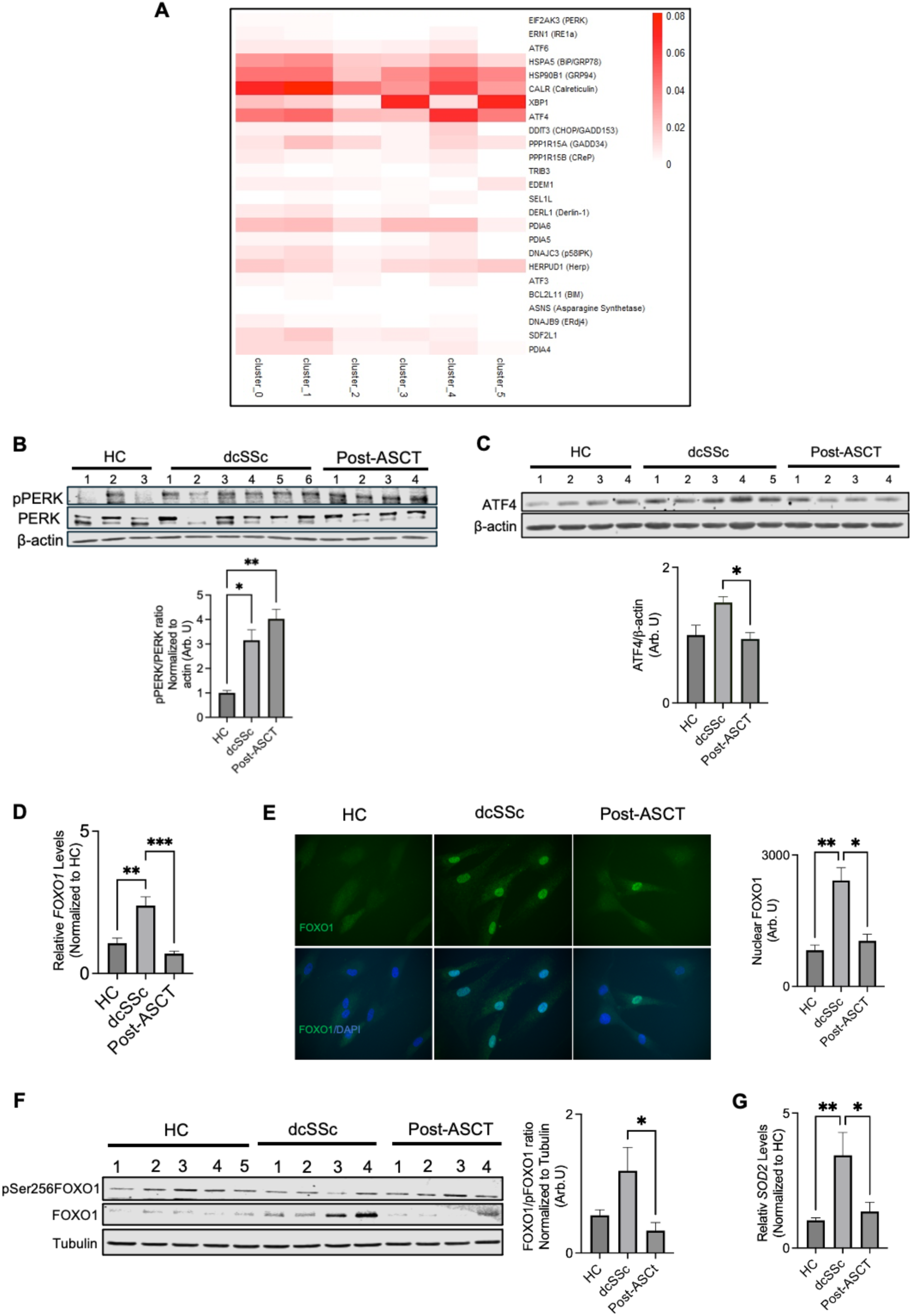
dcSSc FB have increased PERK, ATF4 and FOXO1 activation. **A)** We reanalyzed spatial RNA sequencing data from our recent publication ^47^ and focused on transcripts that are altered in the unfolded protein response. **B)** Relative PERK phosphorylation (activation) and downstream ATF4 translation **C**) was determined using IB and FB from independent biological replicates of HC (n=4), dcSSc (n=5), or post-ASCT (n=4), then plotted the relative frequency of pPERK/PERK or ATF4 levels normalized to beta-actin. **D**) *FOXO1* mRNA levels were determined via qRT-PCR in FB from independent biological replicates of HC (n=5), dcSSc (n=7) and post-ASCT (n=5). **E)** Representative fluorescence microscope image (60x objective) of FOXO1 (green) and nucleus (DAPI, blue). FOXO1 activation was determined by either quantifying the relative fluorescence intensity of nuclear FOXO1 in independent biological replicates of primary FB from HC (n=6), dcSSc (n=7), and post-ASCT (n=5); **F)** or by quantifying the abundance of FOXO1/p-Ser257-FOXO1 (normalized to tubulin as a loading control) in independent biological replicates of primary FB from HC (n=6), dcSSc (n=8), and post-ASCT (n=6). **G)** We confirmed FOXO1 activation by quantifying the mRNA levels of its downstream target *SOD2* in independent biological replicates of primary FB from HC (n=5), dcSSc (n=8) and post-ASCT (n=6). All data were presented as the means ± SEM. Statistical significance was determined by one -way ANOVA. Using GraphPad Prism, statistical significance was determined when *p<0.05, **p<0.005 and ***p<0.0005.

To link genotoxic stress to ATF4/FOXO1 activation, we treated HC FB with etoposide, a known genotoxic agent that induces DSBs, and observed induction of both ATF4 and FOXO1 (**Supplementary fig. 3**). Finally, our co-immunoprecipitation (co-IP) data suggested a direct interaction between PERK and potentially ATF4 (as previously suggested ^50^). FOXO1 co-immunoprecipitated with PERK and ATF4 in dcSSc but not HC FB (**Supplementary fig. 5**).

Together, these findings show that dcSSc fibroblasts, unlike HC or post-ASCT FB, exhibit metabolic adaptations driven by coordinated activation of the PERK/ATF4/FOXO1 axis. *PERK and FOXO1 are required for dcSSc FB survival* We next inhibited PERK or FOXO1 activation using specific PERK ^51^ and FOXO1 ^52^ pharmacological inhibitors. To determine the optimal inhibitor dosing, fibroblasts were treated with a range of concentrations and exposure times. Based on these experiments and results from previous studies ^53–55^, 1 µM for 2 h was selected for the PERK inhibitor and 1 µM for 4 h for the FOXO1 inhibitor. Suppression of either pathway rapidly induced mitochondrial-dependent apoptosis in dcSSc FB, as suggested by increased levels of cCasp 3 and cCasp 9 (**Fig. 6A and B**), whereas HC FB were relatively unaffected (**Fig. 6C, D**). These findings indicate that sustained PERK and FOXO1 activity are required to maintain the survival program preferentially present in dcSSc FB.

**Fig. 6.**
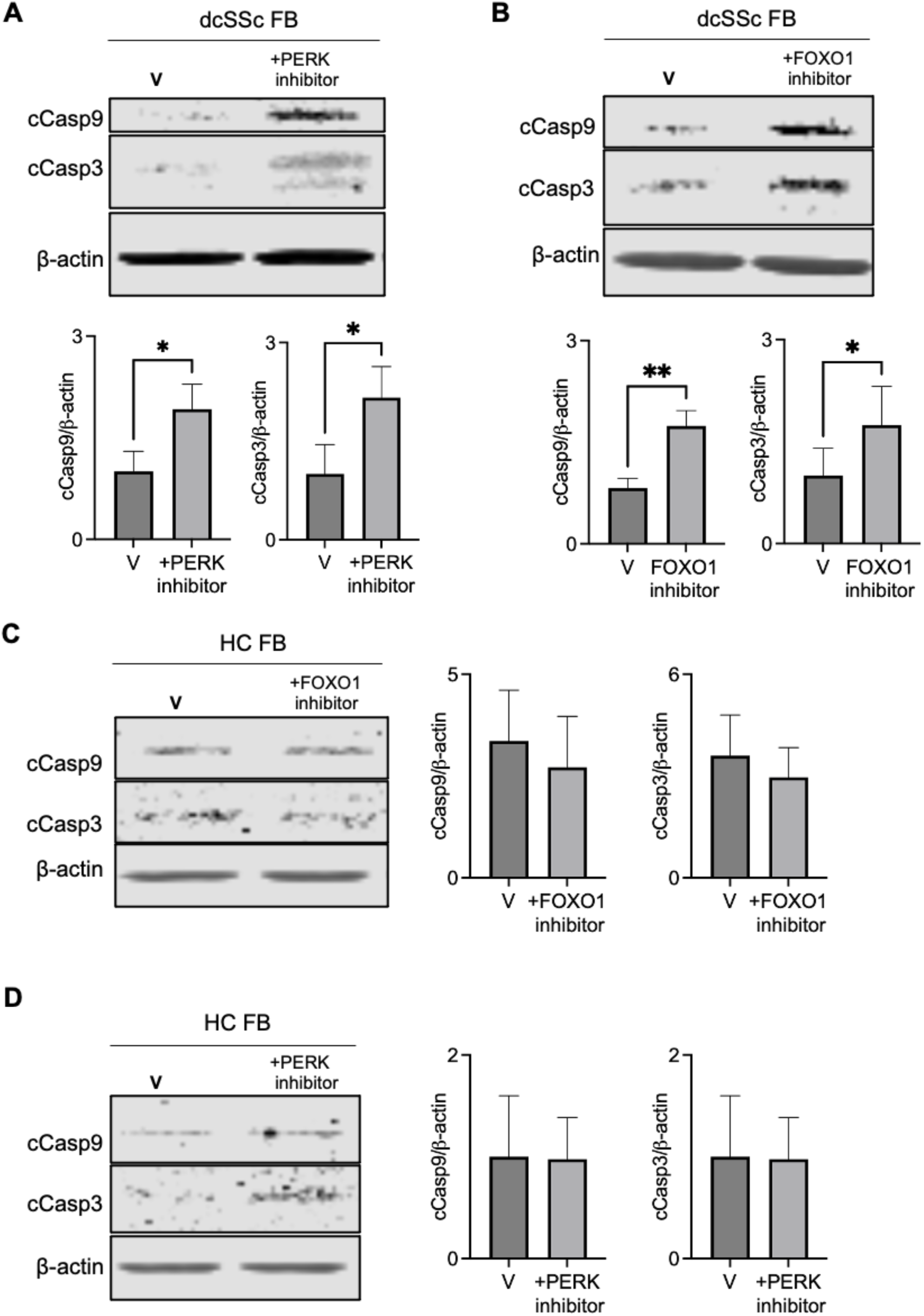
dcSSc require PERK and FOXO1 for mitochondrial dependent survival. dcSSc FB (n=4 X 2 independent biological replicates) were treated with either a **A)** PERK kinase pharmacological inhibitor (GSK 2606414) for 2 hours or **B)** FOXO1 pharmacological inhibitor AS1842856 for 4 hours, then the frequency of cleaved Caspase 3 and 9 were quantified via IB. Treatment of primary FB from HC (n=3 X 2 independent biological replicates) with **(C, D)** either inhibitor does not increase apoptosis as quantified by cleaved Caspase 3 and 9 levels. All data were presented as the means ± SEM. Statistical significance was determined by paired t-test. Using GraphPad Prism, statistical significance was determined when *p<0.05, **p<0.005 and ***p<0.0005.

## Discussion

This study indicates that dcSSc FB evade apoptosis through metabolic stress remodelling driven by activation of PERK/ATF4/FOXO1 axis.

Previous studies have implicated both PERK and FOXO1 in diverse fibrotic pathways. PERK-mediated activation of ATF4 promotes metabolic adaptations that contribute to patients with asbestos-induced lung fibrosis, and pharmacologic PERK inhibition attenuates this fibrotic process in rodents ^56^. Further to this, targeting the PERK/ATF4 axis via agents such as 4-phenylbutyrate may attenuate renal fibrotic reactions ^57^. Similarly, PERK is required for bleomycin-induced lung injury, and genetic deletion of PERK prevents the development of bleomycin-related pulmonary fibrosis in mice ^58^. Furthermore, PERK also acts as a survival signal in myofibroblasts through its interaction with the stimulator of interferon genes (STING)^59^. Notably, STING activation has been associated with senescence-like features in SSc ^60^ and linked to chromosomal instability in this disease ^61^. These findings are consistent with our results and highlight the importance of PERK in promoting metabolic adaptations in dcSSc FB.

Importantly, in FB from patients treated with ASCT, the genotoxic stress burden, apoptotic resistance, and metabolic remodelling were markedly reduced, and the ATF4/FOXO1 axis was no longer activated despite ongoing PERK phosphorylation. The mechanisms underlying these restorative effects of ASCT remain incompletely understood. One possibility is that PERK activation in dcSSc FB is driven by multiple upstream stressors only a subset of these signals may persist following ASCT - although these signals may be insufficient to trigger downstream canonical FOXO1/ATF4 activation. An additional explanation is that ASCT facilitates the differentiation and engraftment of new, non-senescent FB from bone marrow–derived progenitors, gradually replacing the highly stressed, mutation-bearing FB populations characteristic of dcSSc skin. Further to this, ASCT conditioning protocols utilize large doses of 4-HC (approximately 200 mg/kg), which are likely lethal to dcSSc that adapt metabolic reprogramming. ASCT, in turn, may promote the reconstitution of FB that do not harbour genotoxic stress signals associated with senescence.

Although our mechanistic study is highly novel, it has several limitations. The sample size of patient-derived FB was relatively small. However, other mechanistic studies in SSc ^62^ have utilized similar unique patient-derived FB. Our study highlights a role for ASCT in reducing (and potentially normalizing) genotoxic stressors and metabolic reprogramming. Previous studies have suggested that genotoxic stress signals may not be restricted to fibrotic tissue but may also be present in immune cells ^63^. Future studies using matched pre- and post-ASCT tissues from the same individual (with serial skin biopsies and potentially blood samples collected at different times post-ASCT) will be important to further delineate this.

In conclusion, our study proposes a mechanistic explanation for the persistence of FB in dcSSc patients despite this substantial burden of genomic mutations and DNA damage, identifying the PERK/ATF4/FOXO1 axis as a key survival pathway (see *Graphical abstract*). We propose that targeting this axis may offer a novel therapeutic strategy to eliminate pathogenic FB and interrupt fibrosis in dcSSc. Notably, PERK inhibitors are already under investigation as complementary agents for targeting relapsed/refractory acute myeloid leukemia cells ^64^, and FOXO1 targeting has shown efficacy in preclinical diabetic wound-healing models ^65,66^, highlighting the translational potential of this novel pathway for future therapeutic development in systemic sclerosis.

## Acknowledgements

This study was supported by research grants from the Canadian Institutes of Health and Research (CIHR, Grant number 496901 to MO and RG), the National Scleroderma Foundation (to MO) and GlycoNET (to MO and LW). MO is supported by an Arthritis Society IMHA/STAR Award (Award 00049) and LK is supported by a Department of Medicine Ballermann Translational Award and WCHRI Graduate Studentship Award. We would also like to acknowledge the flow cytometry and confocal microscopy core facilities at the University of Alberta for their support.

## Conflicts of interest

There are no conflicts of interest related to this study. R.G. reports carrying out clinical trials for Boehringer Ingelheim and Janssen and has received honoraria as a consultant and/or speaker from AbbVie, Boehringer Ingelheim, Eli Lilly, Janssen, Therakos, Kyowa Kirin, Recordati and Sanofi. M.O. has received honoraria as a consultant and/or speaker for Boehringer Ingelheim and Therakos Inc. M.O. reports carrying out clinical trials for Merck and AstraZeneca. U.M. small shareholder of TSVascular and umcg spin-off company for the development of sodium thiosulphate in clinical medicine. The remaining authors declare no conflicts of interest.

## Data availability

The raw data generated in this study are available from the corresponding author on request.

## Author contributions

LK: writing first and editing final draft, conceptualization, design, analysis of data, data interpretation, and performing experiments.

JW: editing final draft, conceptualization, design, analysis of data, data interpretation, and performing experiments.

CH: editing final draft, analysis of data, data interpretation, performing experiments

LW: editing final draft, supervision and resources

CvE: editing final draft, analysis of data

DR: editing final draft, resources

CD: editing final draft, resources

JS: editing final draft, resources

KJ: editing final draft, resources

UM: editing final draft, resources

EN: editing final draft and resources

MB: editing final draft and resources

JP: editing final draft, resources

JWCT: editing final draft, supervision and resources

HV: editing final draft, supervision and resources

RG: writing first and editing final draft, conceptualization, design, analysis of data, data interpretation, supervision, funding acquisition.

MO: writing first and editing final draft, conceptualization, design, analysis of data, data interpretation, supervision, funding acquisition.

## Supporting information

**Supplementary table 1:** Patient demographics and clinical information.

|  | dcSSc | Post-ASCT |  |
| --- | --- | --- | --- |
| Categorical variables | Count (%) | Count (%) | p-value |
| Sex (F) | 5/10 (50.0) | 3/8 (37.5) | 0.596 |
| ILD (Y) | 4/10 (40) | 4/8 (50) | 0.671 |
| Anti-scl70 Ab (Y) | 3/10 (30) | 4/8 (50) | 0.387 |
| Anti-RNA polymerase III Ab (Y) | 6/10 (60) | 3/8 (37.5) | 0.343 |
| Antinuclear Ab (Y) | 10/10 (100) | 8/8 (100) | - |
| Medications |  |  |  |
| None | 1/10 | 5/8 |  |
| Vasodilators | 0/10 | 3/8 |  |
| Immunomodulatory therapies(IS) | 7/10 | 0/8 |  |
| Combined Vasodilators and IS therapies | 2/10 | 0/8 | 0.002 |
| Continuous variables | Median (IQR) | Median (IQR) | p-value |
| Age (yrs) | 46.5 (40;54) | 44.5 (40;47.5) | 0.499 |
| mRSS | 21.5 (17;32) | 12.5 (9.5;14.5) | 0.011 |
| Disease duration (pre-ASCT) (yrs) | 1.45 (1.1;2.4) | 3.45 (2.85;4.25) | 0.001 |
| Duration (post ASCT) (yrs) | - | 1.1 (1;2.5) | - |

F-female; Y=yes; Ab-antibody; yrs-years; IQR-interquartile range

**Supplementary fig. 1.**
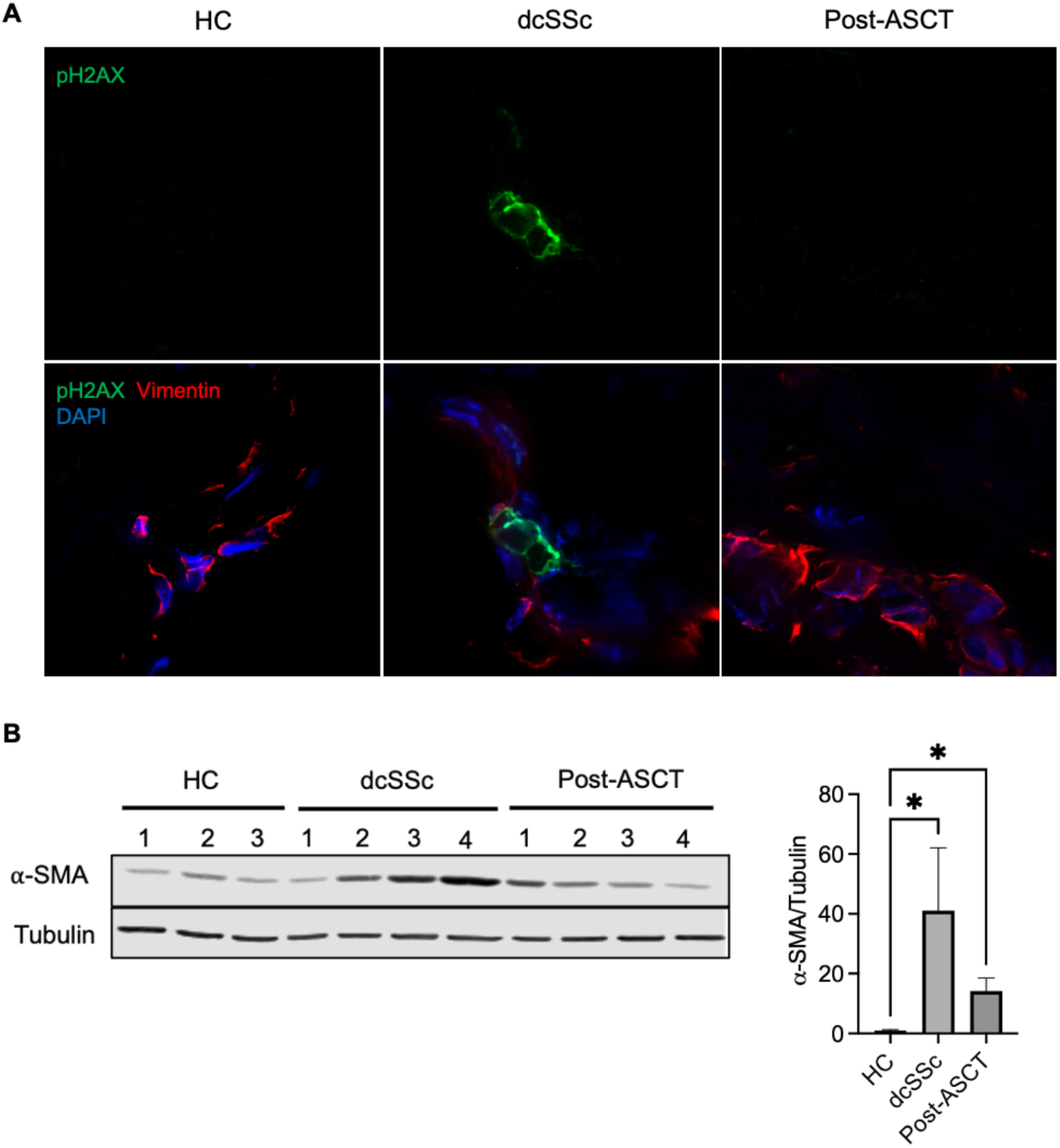
Representative images of double-stranded DNA breaks in skin sections and higher ⍺-SMA in fibroblasts from dcSSc. A) Confocal immunofluorescence images of pH2AX (green) and DAPI nuclei (blue) at 60X in skin sections of HC, dcSSc, and post-ASCT to analyze DNA damage foci. B) Representative IB of ⍺-SMA protein in FB from different groups and tubulin was used as a loading control (n: HC=4, dSSc=8 and post-ASCT=7, independent biological replicates).

**Supplementary fig. 2.**
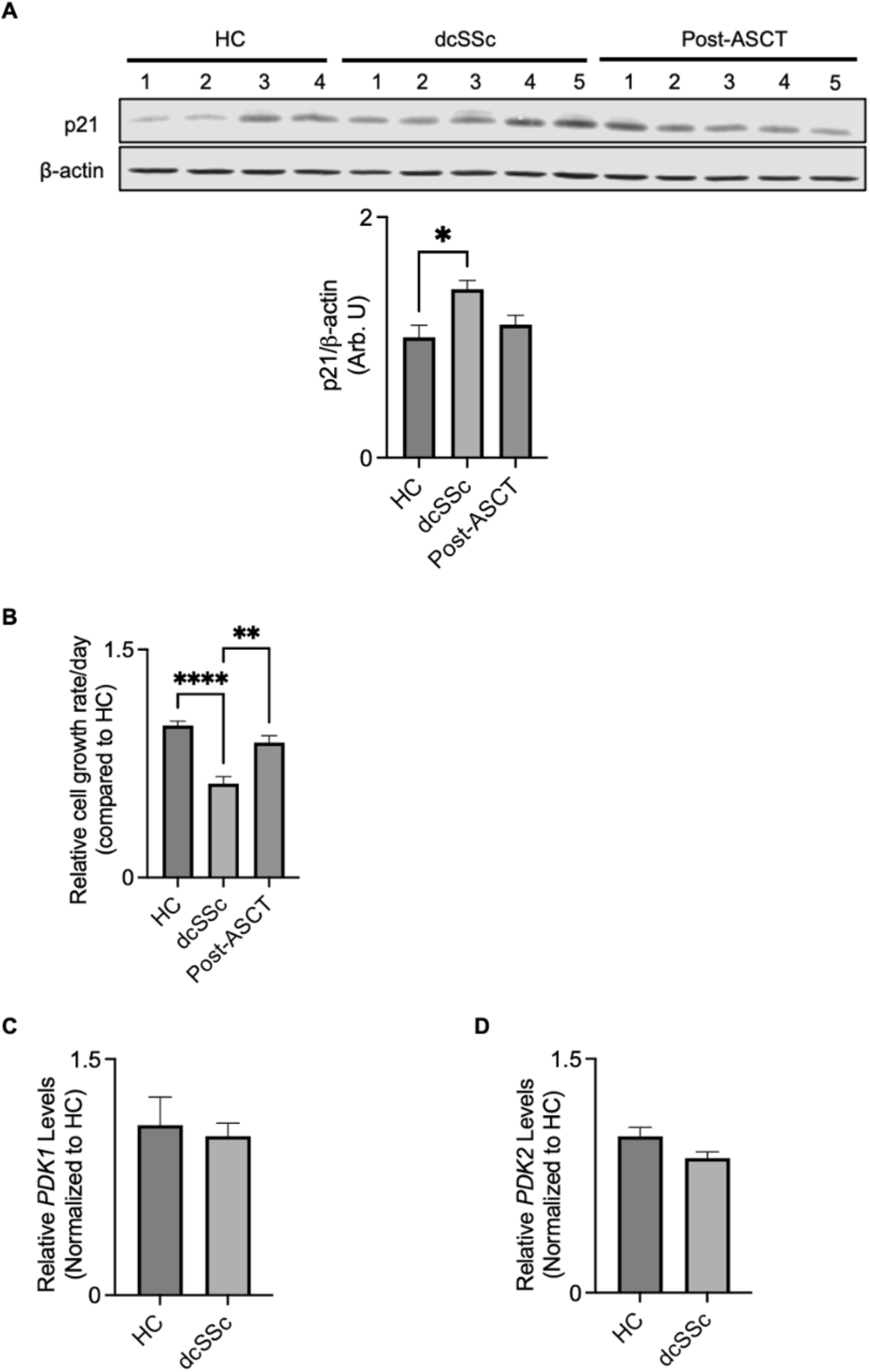
Senescence marker p21 increases and growth rate decreases in dcSSc FB with no change in pyruvate dehydrogenase kinase (PDKs), PDK1, PDK2. A) Immunoblot analysis of senescence-associated signals p21 (normalized to ß-actin) upregulated in dcSSc FB than HC and post-ASCT (N: HC=5, dcSSc=8 and post-ASCT=6). B) After seeding equal numbers of cells, FB growth was quantified by cell counts, revealing a reduced growth rate in dcSSc FB compared with HC and post-ASCT. (N: HC=5, dcSSc=5 and post-ASCT=5, independent biological replicates). C) PDK1 and D) PDK2 mRNA levels are not different between HC and dcSSc FB as determined by qRT-PCR analysis.

**Supplementary fig. 3.**
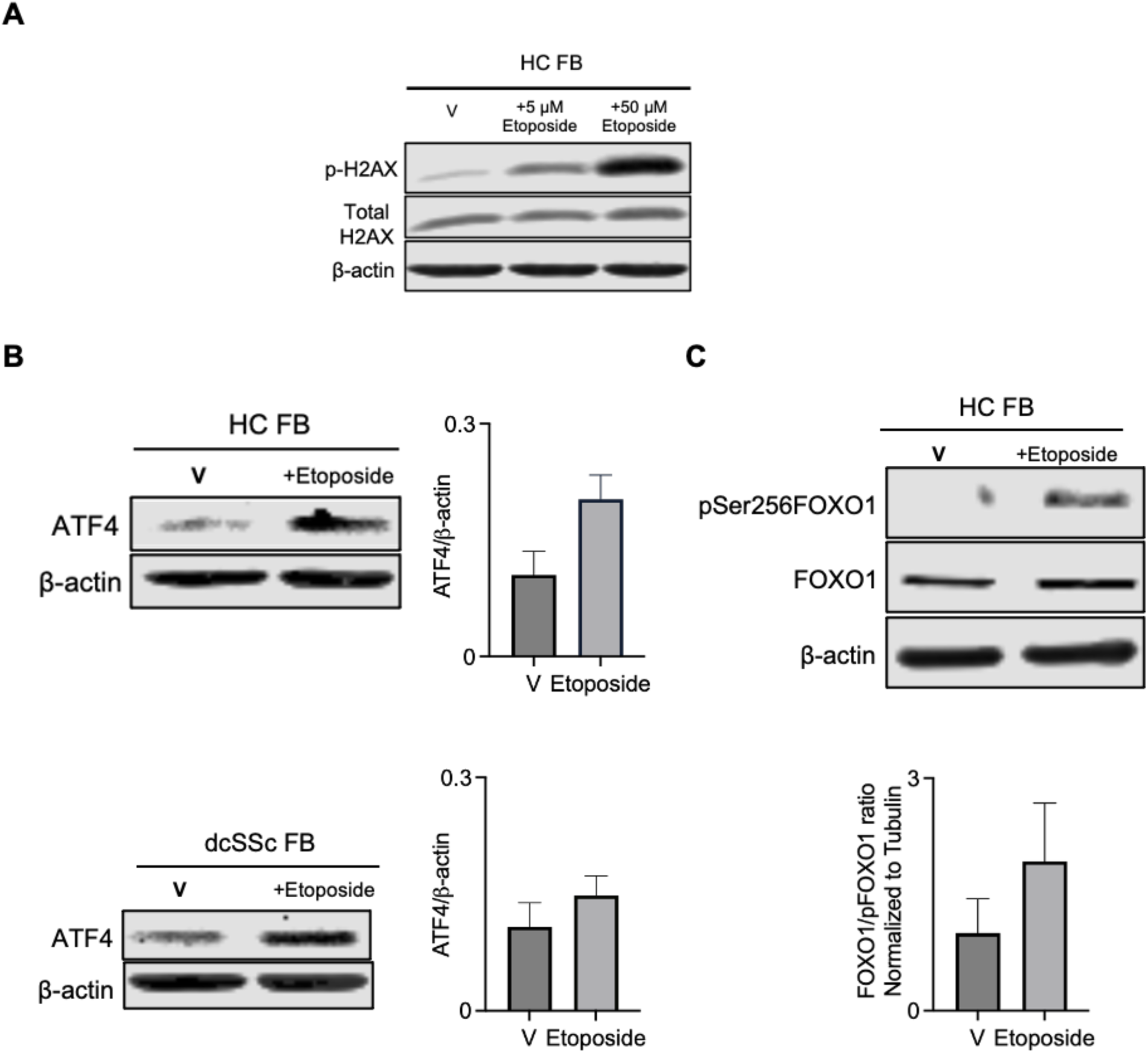
DNA damage directly promotes ATF4 and FOXO1 activation. FB from HC or dcSSc were treated with etoposide (50 µm) 2 hours, then A) p-H2AX, ATF4 levels (B), or FOXO1 activation (via Ser256 phosphorylation) (C) was assessed via IB. Data representative of 2 independent experiments.

**Supplementary fig. 4.**
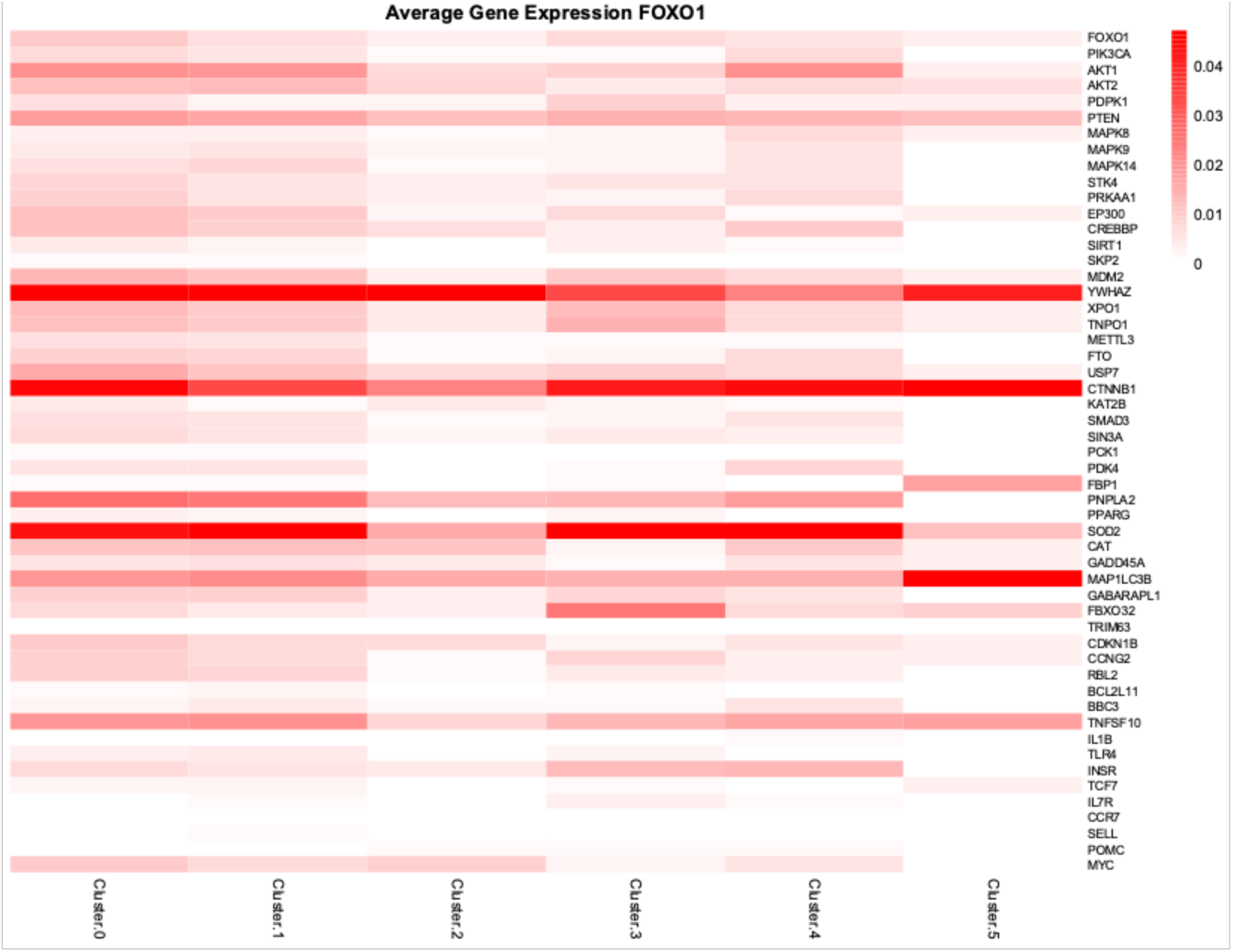
SSc dermal sections have increased expression of known FOXO1 targets in fibroblast clusters (clusters 0 and 1). Spatial RNA sequences were re-analyzed from our recent publication ^47^. Relative abundance of transcripts that are downstream of FOXO1 in each cluster were determined, as we previously described ^47^.

**Supplementary fig. 5.**
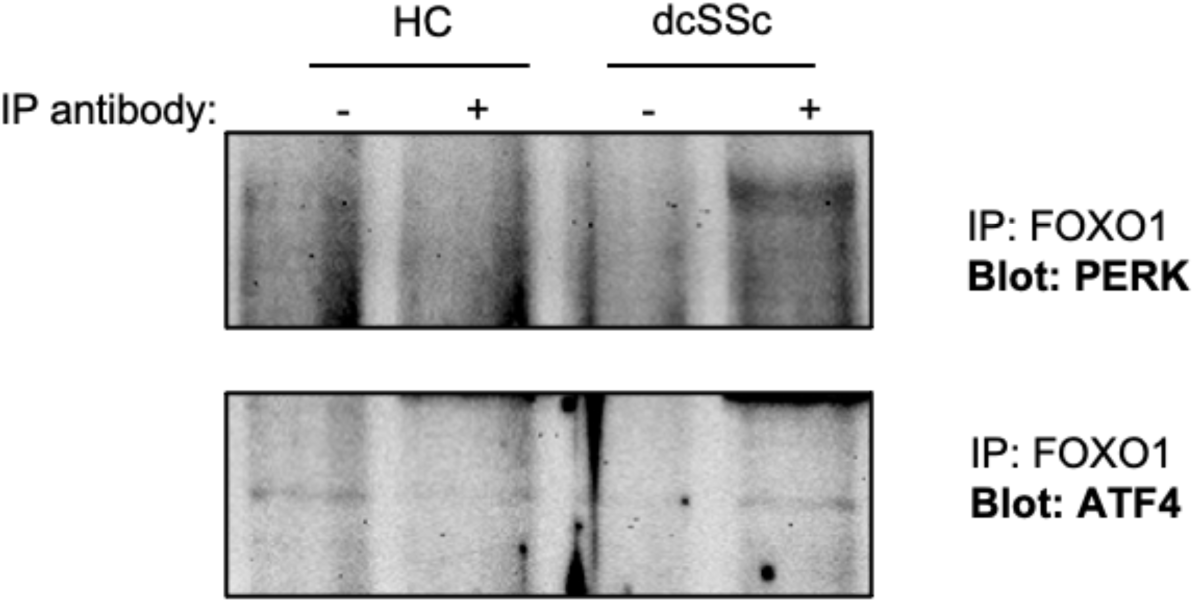
FOXO1 directly associates with PERK and ATF4. Primary FB lysates from HC, or dcSSc were used for co-immunoprecipitation with FOXO1 (capture antibody) then the co-immunoprecipitated was resolved used SDS/PAGE and either PERK or ATF4 association was determined via IB. Representative blots from 2 independent experiments.

## Notes

### Competing Interest Statement

The authors have declared no competing interest.

